# Interwoven DNA-MOF crystals for multi-millennial data storage

**DOI:** 10.1101/2025.07.03.662672

**Authors:** Bingyi Liu, Zhishang Li, Zimu Li, Mingqiang Li, Yang Liu, Zhongyu Wei, Qien Shi, Liqi Zhou, Huating Kong, Kai Xia, Haitao Song, Xiaoguo Liu, Chunhai Fan, Fei Wang

**Affiliations:** State Key Laboratory of Synergistic Chem-Bio Synthesis, School of Chemistry and Chemical Engineering, Frontiers Science Center for Transformative Molecules, Institute of Translational Medicine, Shanghai Jiao Tong University, Shanghai 200240, China; Institute of Materiobiology, College of Sciences, Shanghai University, Shanghai 200444, China; Shanghai Synchrotron Radiation Facility (SSRF), Shanghai Advanced Research Institute, Chinese Academy of Sciences, Shanghai 201204, China; Jiaxing Key Laboratory of Biosemiconductors; Xiangfu Laboratory, Jiashan, Zhejiang 314102, China; The Institute of Artificial Intelligence and National Center for Translational Medicine, Shanghai Jiao Tong University, Shanghai 200240, China; Institute of Molecular Medicine, Shanghai Key Laboratory for Nucleic Acids Chemistry and Nanomedicine, Renji Hospital, School of Medicine, Shanghai Jiao Tong University, Shanghai 200127, China

## Abstract

DNA has emerged as a promising medium for high-density and energy-efficient data storage, providing a promising solution to capacity constraints and energy demands resulting from the exponential growth of data. Long-term stable storage of DNA data in ambient conditions is a critical requirement for its practical application. Whereas strategies have been proposed to protect DNA from environmental degradation, they are often at the expense of data storage density. Here, we propose a strategy via interweaving metal-organic framework (MOF) and DNA to synthesize DNA-MOF crystal (DNA-MOF), a high-density and long-lifespan DNA-based information material. The uniform distribution of DNA molecules within the crystalline frameworks enables a high loading capacity of up to 30 wt%, corresponding to a theoretical information density of 106.6 exabytes/g. The crystal structure of MOF-DNA demonstrates efficient resistance to environmental damage such as heat, ultraviolet radiation, DNase, etc., achieving an estimated archival half-life over 24,000 years at 9.4 °C. The interwoven DNA-MOF crystals establish a material foundation for the reliable operation of DNA storage systems in open-field deployment and provide a new pathway toward sustainable long-term management of massive data.

## Introduction

Currently, human-generated data is growing exponentially, with global total projected to reach 394 ZB by 2028^1^. Due to the capacity limitations and energy demands of current storage media, archiving and managing such vast amounts of data is becoming increasingly unfeasible. DNA offers exceptional potential for long-term data storage, owing to its ultra-high theoretical information density (∼455 EB/g), remarkable durability, and minimal energy requirements for data maintenance^2^. Although DNA exhibits remarkable molecular stability—evidenced by accurate gene recovery from million-year-old fossils^3,4^ —certain common environmental factors (such as heat, enzyme, ultraviolet radiation, oxidation) may lead to degradation of DNA and consequent data loss, posing a critical challenge for practical data storage applications^5,6^. Therefore, shielding DNA from such environmental damage is essential to realize its inherent advantages and to enable the transition from laboratory settings to real-world, large-scale applications of DNA storage.

Several strategies including co-precipitation, adsorption, and core-shell encapsulation, have been proposed to protect DNA from degradation^7,8^. The co-precipitation strategy precipitate DNA from solution with inorganic salts such as calcium phosphate, forming hybrid aggregates in which DNA is embedded within the precipitated matrix to enhance stability^9,10^. The adsorption strategy leverages chemical or electrostatic interactions to immobilize DNA onto or into particles, exploiting steric hindrance to resist against enzymatic digestion, thermal degradation, and other stressors^11–13^. Core-shell encapsulation employs materials such as amorphous silica to enclose DNA within sealed structures that offer effective shielding from environmental factors, enabling information preservation for up to 2,000 years at 4 °C^14–16^. Crucially, these methods exhibit fundamental trade-offs between storage density and stability. Co-precipitation can offer relatively high DNA loading (∼20 wt%), but the resulting material often suffers from poor control over component composition, shell thickness, and structural uniformity, limiting its effectiveness in isolating DNA from the environment. The adsorption-based approach generally involves relatively weak interactions, which limits both the loading capacity and the overall protective performance. In contrast, core-shell structures can ensure millennia-scale stability, but only the outer shell is used for DNA storage, resulting in low spatial utilization and limited data density. Therefore, fully exploiting the internal space of protective materials is key to achieving high-density DNA information storage while meeting the requirements for long-term data retention.

Metal-organic framework (MOF) materials are highly crystalline porous materials with periodic and tunable frameworks. Owing to their low density, high stability, and tunable morphology, MOFs have been widely employed for loading organic small molecules, proteins, and DNA in applications including drug delivery^17–19^, catalytic bioreactors^20–22^, as well as molecular separation and recognition ^23–25^. The coordination-bonded crystalline frameworks of MOFs confer intrinsic thermal resilience while tunable pore metrics synergize with engineered hydrophobicity to shield encapsulated payloads. These dual-attribute protection mechanisms position MOFs as promising materials for high-density, long-term DNA data storage. Despite pioneering demonstrations of MOF-mediated DNA preservation^26^, in these approaches DNA is either adsorbed or coordinated onto the surface of preformed MOF crystals^27–29^, or adsorbed onto other particles and subsequently encapsulated by MOFs^30^. In both cases, the internal space provided by MOFs are not fully utilized, resulting in a DNA loading capacity that remains below 5 wt%.

Here, we report a strategy for interweaving DNA with MOFs to synthesize DNA-MOF crystals, in which DNA strands are uniformly and densely filled in the internal space of MOF particles, enabling dense digital data retention. During the crystallization process, the information-encoded DNA molecules serve as heterogeneous nucleation templates, heterocoordinate with metal nodes, and guide the formation of a crystalline structure analogous to Zeolitic Imidazolate Framework-8 (ZIF-8, Fig. 1a). The stored information can be rapidly and nearly lossless retrieved through crystal dissociation using metal chelators, realizing high-fidelity data storage. The interweaving of DNA and MOFs maximizes utilization of the vacancies, enabling a DNA loading capacity of up to 30 wt% (Fig. 1b). Meanwhile, the protective properties of the MOF crystals against environmental stressors such as UV irradiation, heat, and enzymatic degradation are preserved in DNA-MOFs. By employing DNA-MOFs, digital information can be stored at a physical density of 106.6 EB/g for over 24,000 years at 9.4 °C (Fig. 1c), enhancing the practicality and sustainability of DNA-based data storage, and providing a viable medium for the long-term preservation of massive data.

**Fig. 1.**
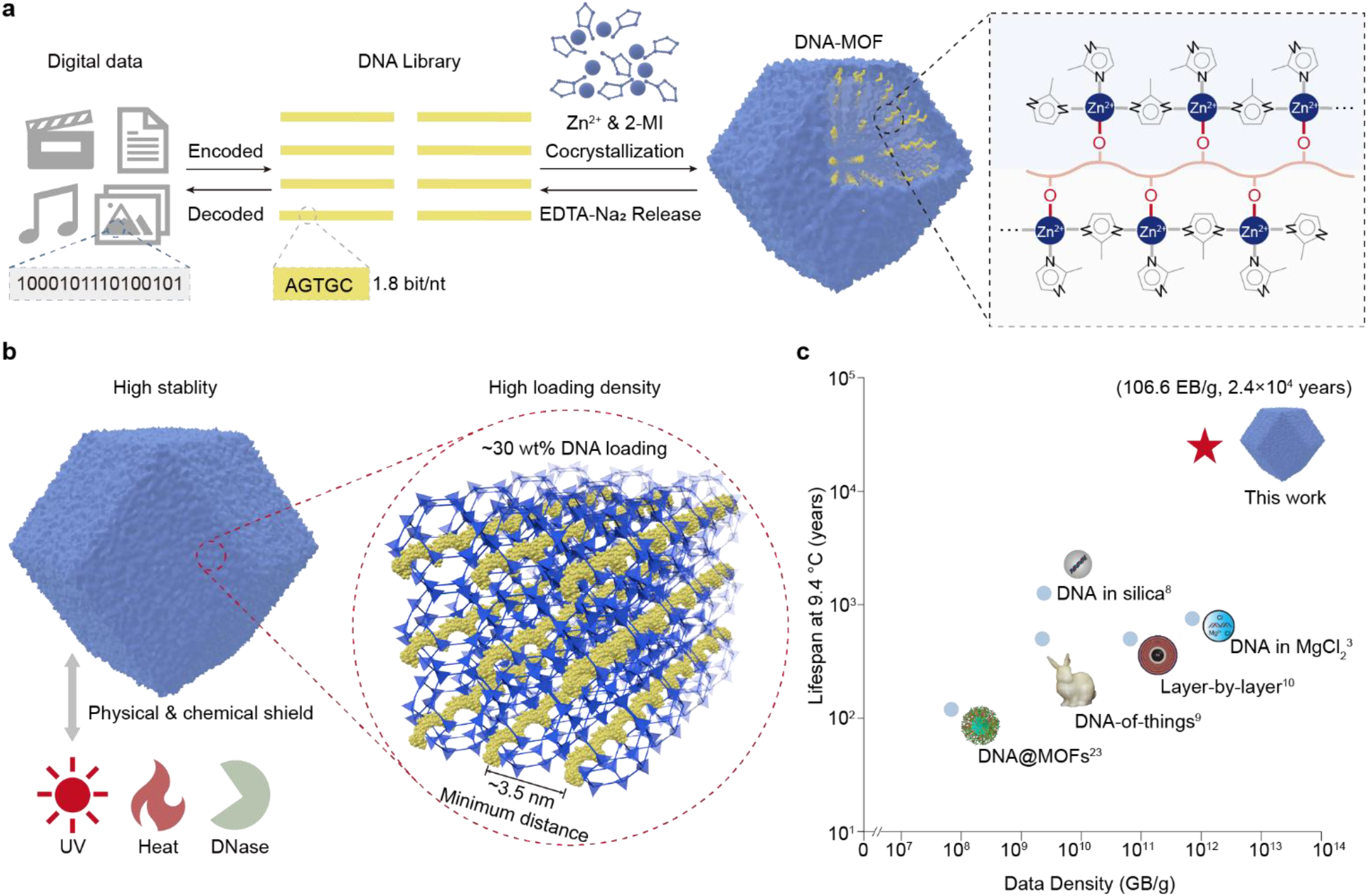
Overview of high-density, long-term DNA storage enabled by DNA-MOFs. (a) Workflow illustrating file-encoded DNA interwoven DNA with MOF via heterocoordination to form DNA-MOF crystals for long-term data preservation. (b)The DNA strands interweaving with ZIF-8 frameworks by Zn-O coordination, densely filled within the crystal lattice to achieve high information storage density, while the crystalline structure protects the DNA during preservation. (c) A comparison to prior work shows that DNA-MOF simultaneously provides both high storage density and long lifespan.

## Results

### Synthesis and characterization of interwoven DNA-MOF crystals

To construct a high-density DNA storage system resistant to environmental interference, we engineered an interweaving MOF-DNA architecture where heterocoordination between DNA strands and ZIF-8 metal nodes achieves uniform distribution of DNA throughout the internal vacancies of the crystals (Fig. 2a). We selected ZIF-8, one of the representative MOF structures with excellent hydrothermally stability and biomolecular compatibility, as the target framework. Since Zn^2+^ can interact with DNA through both electrostatic interaction and coordination, premature binding between Zn^2+^ and DNA during the early stages of crystallization may lead to conformational changes and aggregation of the DNA strands^31,32^, thereby hindering subsequent coordination of Zn^2+^ and disrupting the ordered growth of the crystal. To address this, we first mixed DNA with 2-methylimidazole in aqueous solution, followed by the addition of Zn^2+^, thus guiding the interweaving of DNA with MOF during the nucleation and growth processes.

**Fig. 2.**
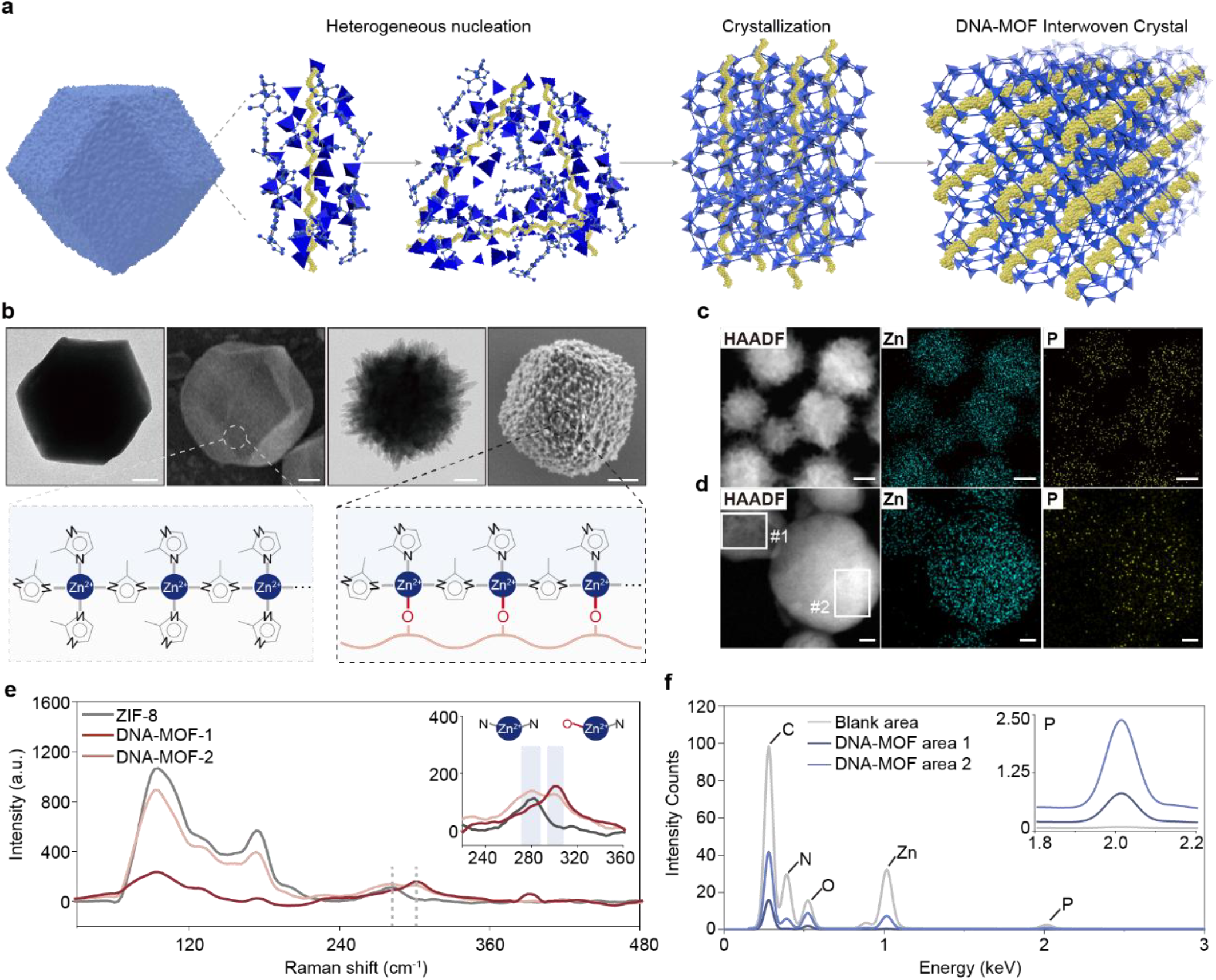
Validation of DNA-MOF interweaving strategy. (a) Schematic diagram of the proposed interweaving of MOF and DNA. During the crystallization process, Zn^2+^ ions accumulate around the phosphate groups of the DNA backbone, promoting crystal growth through a heterogeneous nucleation mechanism. (b) Representative TEM and SEM images of ZIF-8 (left) and DNA-MOF (right). Scale bars: 100 nm (left), 50 nm (right). (c) HAADF-STEM image and corresponding EDS elemental maps of Zn and P of DNA-MOF. Scale bar: 100 nm. (d) HAADF-STEM image and corresponding EDS elemental maps of Zn and P in ultrathin sections of DNA-MOF. Scale bar: 50 nm. (e) Raman spectra of ZIF-8 and DNA-MOFs with two different particle sizes. (f) EDS spectrum of the framed areas in DNA-MOF ultrathin sections. The inset spectrum highlights the characteristic peaks of P element.

We observed that DNA-MOF particles showed distinct morphological features differing from conventional ZIF-8 crystals, exhibiting rough surfaces with spiky protrusions (Fig. 2b). These unique morphological characteristics may originate from lattice defects introduced by DNA participation in the crystallization process through heterocoordination with Zn^2+^. In contrast to the homogeneous nucleation process of ZIF-8 in solution, DNA molecules can serve as heterogeneous nucleation templates by enriching Zn^2+^, thereby reducing the free energy barrier for nucleation and accelerating the local nucleation process on DNA interfaces^33^.. We employed Raman spectroscopy to verify the coordination of DNA within the crystal framework. Compared to the Raman spectrum of ZIF-8, the DNA-MOF spectrum displayed a distinct peak at ∼302 cm^-1^ (Fig. 2c), which appeared as shoulder adjacent to the characteristic Zn-N stretching vibration at ∼282 cm^-1^ of ZIF-8^34,35^. This confirms that DNA altered the coordination environment of Zn^2+^, demonstrating coordination between Zn^2+^ and phosphate O atoms from DNA backbone. The distinct Raman features suggest that DNA is not merely physically encapsulated by the MOF, but rather incorporated as a structural component during the crystallization process.

### Optimization of data loading in DNA-MOFs

To evaluate DNA-loading capacity, DNA-MOF particles were dissolved to release the DNA strands. The DNA loading percentage (wt%) was calculated by measuring the mass of both the MOF component and the released DNA (Fig. 3a). Specifically, Zn^2+^ ions in the framework were chelated using 20 mM EDTA-Na^236^, which enabled complete DNA release within 10 minutes (Fig. 3b). Following this, we investigated DNA loading efficiencies under various initial DNA concentrations. The loading percentage correlated positively with DNA concentration, reaching saturation at ∼50 μM (Fig. 3c). The maximum DNA loading achieved was 30 wt%, corresponding to a theoretical data storage density of 106.6 EB/g.

**Fig. 3.**
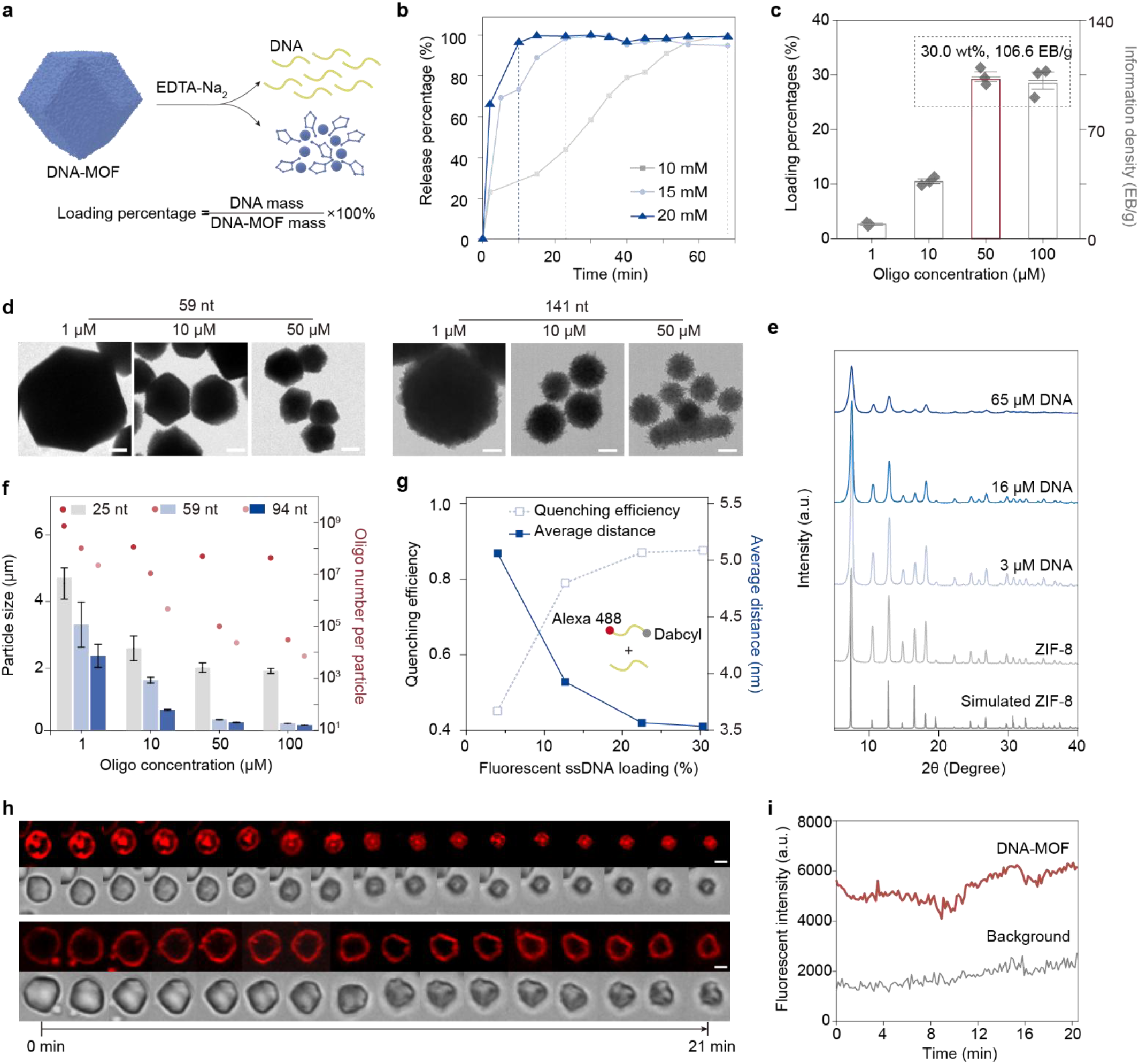
Optimization and characterization of DNA loading in DNA-MOFs. (a) Schematic illustration showing the release of DNA from DNA-MOFs. (b) DNA release kinetics from DNA-MOFs under varying concentrations of EDTA-Na_2_. (c) DNA loading percentages in DNA-MOFs synthesized with varying initial oligo concentrations. (d) Representative TEM images of DNA-MOFs synthesized with different oligo lengths and concentrations. (e) PXRD patterns of ZIF-8 and DNA-MOFs synthesized with varying DNA concentrations. (f) Particle sizes of DNA-MOFs with different oligo lengths and concentrations and calculated number of oligos per particle. (g) Quenching efficiency and corresponding averaged fluorophore-quencher distance within DNA-MOFs. (h) Confocal images of representative DNA-MOF particles during the release process of Alexa647-labeled DNA. Scale bar: 2 μm. (i) Fluorescence intensity variation along the contour of a representative particle over time.

The DNA-MOF interweaving strategy exhibited broad applicability across oligonucleotide strand lengths (25-8064 nt,). Saturation loading efficiencies remained consistently high (20–30 wt%), confirming that DNA-MOF enables efficient and sequence-length-independent DNA encapsulation and protection. At maximum loading of 94 nt oligos, each DNA-MOF particle was calculated to contain ∼7700 oligos (Fig. 3f), equating to a theoretical maximum data storage density of ∼159 KB per particle.

To probe the spatial organization of ultrahigh-loaded DNA within the DNA-MOF crystals, powder X-ray diffraction (PXRD) was performed with varying loading levels. Compared with ZIF-8, DNA-MOFs exhibited progressively broadened diffraction peaks, decreased intensities, and partial peak disappearance as DNA loading increased (Fig. 3e). These observations suggested that higher DNA incorporation introduces heterocoordination sites, increasing lattice defects and reducing crystallinity. Correspondingly, the particle size decreased with increasing DNA concentration for each length of loaded DNA (Fig. 3d), likely due to defect-induced crystal growth inhibition. Structural integrity was further probed through gas sorption analysis. Although the average pore diameter remained unchanged relative to ZIF-8, the intensity of the pore size distribution decreased and the specific surface area declined by 17.7%. This may be attributed to the aberrant coordination between DNA and Zn^2+^, introducing lattice defects or distortions and resulting in the loss of some pores.

To further examine DNA spatial distribution within the framework, we co-crystallized ssDNA labeled with a fluorophore-quencher pair (Alexa 488/Dabcyl) at each end. Fluorescence quenching increased with DNA loading. This trend confirmed that DNA disperses uniformly within MOF pores without aggregation and the inter-strand quenching plays the dominant role. To exclude the influence of particle size variation, we fixed the total DNA concentration and varied the of fluorescently labeled. As the proportion of labeled DNA increased, quenching efficiency correspondingly rose. Based on quenching efficiency calculations, the average minimum intermolecular distance between DNA strands was ∼3.5 nm (Fig. 3g), consistent with the high loading of DNA within DNA-MOF crystals. In addition, the Alexa 647-labeled DNA-MOF dissolution process was in situ observed by confocal microscopy. We observed that for most particles, prominent fluorescence intensities appeared at the particle periphery during EDTA-induced disassembly (Fig. 3h). Due to limited laser penetration depth and partial reflection of the excitation light at the particle surface, fluorophores near the particle surface at the focal plane are preferentially excited. Quantitative profiling revealed invariant peripheral intensity across reduced particle sizes (Fig. 3i), confirming that DNA is uniformly distributed throughout the DNA-MOF particles and released in a spatially homogeneous manner.

### Environmental resilience of DNA-MOFs

To evaluate the protective capability and lifespan of DNA-MOFs under practical conditions, we exposed DNA-MOFs to environments that cause physical, chemical, and biological damage on naked DNA. Thermogravimetric analysis (TGA) revealed that DNA-MOF exhibited significantly enhanced thermal stability compared to DNA (Fig. 4a). Specifically, DNA-MOF showed negligible weight loss below 200 °C and only gradual mass reduction between 200 °C and 600 °C. In contrast, free DNA displayed marked degradation below 200 °C and underwent rapid decomposition above 230 °C. This enhanced thermal stability is attributed to the SOD-type topology shared between DNA-MOF and ZIF-8. The distinct slow decomposition behavior of DNA-MOF at elevated temperatures may arise from the incorporation of DNA, which disrupts the original periodic Zn-N coordination network by introducing partial Zn-O interactions.

**Fig. 4.**
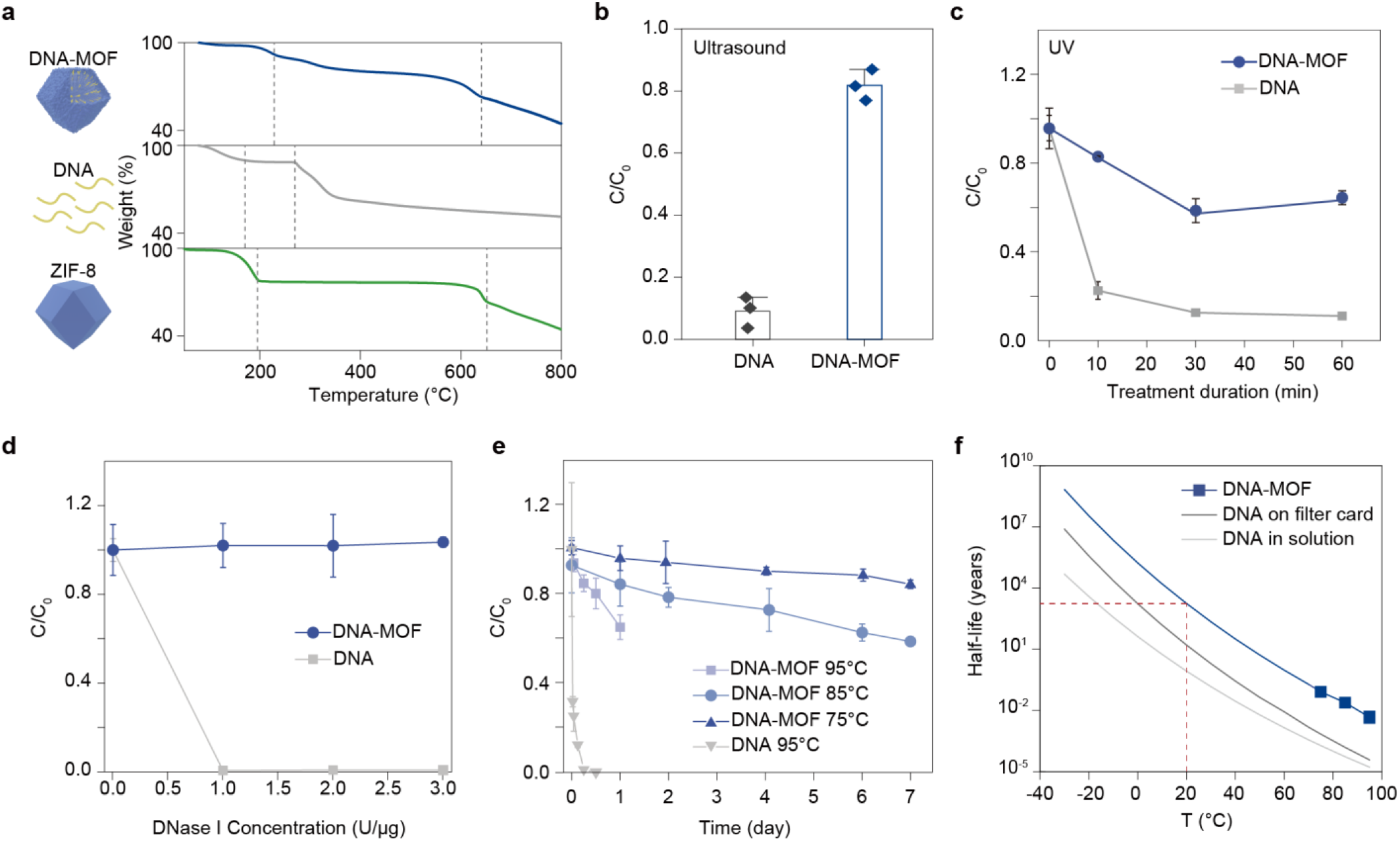
Stability of DNA under degradative environmental conditions. (a) Thermogravimetric analysis of DNA-MOF, DNA and ZIF-8. (b) Relative concentration of DNA remaining after 30 minutes of ultrasonic treatment. (c) Degradation kinetics of unprotected DNA and DNA-MOF under UV exposure. (d) Relative concentration of DNA remaining after 10 minutes of treatment with DNase I at different concentrations. (e) Degradation kinetics of DNA stored in solution versus DNA-MOF under elevated temperatures. (f) Estimated half-life of 94-nt ssDNA stored in DNA-MOF, compared with DNA stored on FTA filter card^14^, in aqueous solution^38^, and in moa bone^39^.

To access mechanical stability, we sonicated free DNA and DNA-MOF solutions for 30 minutes. Quantitative analysis showed that 90.9% of the free DNA degraded, whereas only 18.2% of the DNA released from DNA-MOFs was damaged (Fig. 4b). Ultrasonic degradation mainly results from cavitation-induced shear forces^30^; the rigid framework of DNA-MOF and the homogenous distribution of DNA strands within the crystalline lattice help to dissipate mechanical stress and thus protect DNA from damage. Photostability testing under UV-A irradiation (365 nm) showed 89.0% degradation of free DNA after 1 hour, compared to 35.6% for DNA within DNA-MOFs (Fig. 4c). This protective effect against UV exposure is attributed to the ZIF-like crystal structure of DNA-MOF, which exhibits limited transparency in the UV range and thus provides effective shielding and spatial isolation for the encapsulated DNA.

In enzymatic degradation tests, DNA-MOFs demonstrated substantial resistance against DNase. When treated with 3.0 U/μg DNase I, unprotected DNA showed a degradation rate of 99.6%, whereas only 2.2% of the DNA in DNA-MOFs was degraded (Fig. 4d). This protection is attributed to the sub-nanometer pore sizes of DNA-MOF, which are smaller than the dimensions of DNase I and other common nucleases, thereby creating a physical exclusion barrier by the ordered crystalline structure.

To quantitatively assess the long-term durability of DNA-MOF, we performed accelerated aging experiments. Under 75 °C, 85 °C, and 95 °C, DNA-MOFs retained a significantly higher fraction of intact DNA compared to free ssDNA. For instance, after 1 day at 95 °C, DNA-MOFs retained 65% of released DNA, while free DNA exhibited only 0.41% retention within 0.5 days (Fig. 4e). Fitting the decay rates at different temperatures yielded an Arrhenius-type activation energy (Ea)^14^ of 155.34 kJ/mol for 94-nt ssDNA in DNA-MOFs, which is substantially higher than that of DNA in solution and slightly higher than that of DNA stored on filter paper. Based on this analysis, the estimated half-life of DNA stored in DNA-MOFs is approximately 24,300 years at 9.4 °C and 2,210 years at room temperature (20 °C) (Fig. 4f).

### Information integrity in DNA-MOF

To further evaluate the information preservation integrity of DNA-MOF, we encoded digital information into a DNA library comprising 10,800 distinct strands, loaded the DNA library into DNA-MOF particles, and subsequently released for sequencing (Fig. 5a). To directly display the data preservation capability of DNA-MOF, we employed an encoding method without error-correction. Under this scheme, mutations and strand degradations can be directly mapped to errors in the retrieved digital file. Analysis of the frequency distribution of perfect calls per million reads (PCPM) revealed comparable average PCPM coverage between the original DNA library (65.4) and the DNA-MOF-released library (65.7) (Fig. 5b). Furthermore, error rate distributions closely matched between the two groups, with average deletion and insertion error rate differences <0.002%, and a marginal increase of only ∼0.01% in substitution error rate per sequence in the DNA-MOF sample (Fig. 5c, d)

**Fig. 5.**
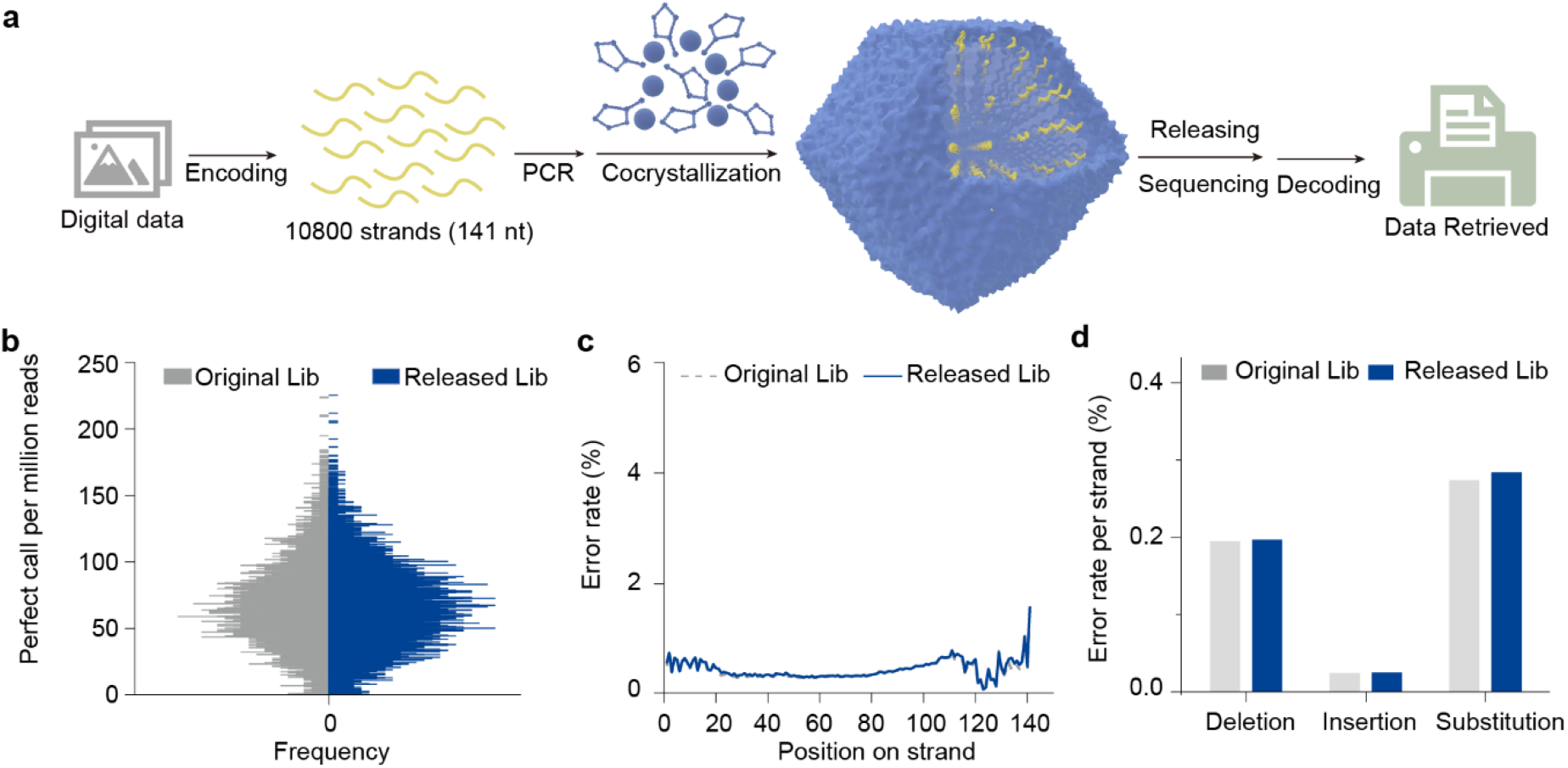
Digital data preservation in DNA-MOFs and information integrity evaluation. (a) Schematic diagram of the DNA-MOF digital data storage workflow. (b) Frequency distribution of perfect calls per million reads for original DNA library and DNA-MOF-released DNA library. (c) The error rates for each DNA sequence position of original DNA library and DNA-MOF-released DNA library. (d) Error type distribution profiles of original DNA library and DNA-MOF-released DNA library.

These results confirm that the processes of crystallization and the release of DNA from DNA-MOFs preserved the integrity of the stored data without introducing significant errors or degradation, thereby. Compared to previously reported protective DNA storage methods that utilize acidic denaturation for DNA retrieval^9^, the use of EDTA-Na_2_ offers a milder alternative that minimizes chemical alterations such as base deamination, thus reducing the risk of information corruption during data extraction.

### Multi-millennial-scale digital data storage within DNA-MOFs

To further evaluate the lifespan of data storage using DNA-MOFs, we conducted accelerated aging experiments by incubating both DNA-MOFs and unprotected DNA libraries at 95 °C. Compared to the unprotected group, the DNA-MOFs demonstrated notable advantages in preserving sequence integrity. During 7 days of 95 °C treatment (equivalent to ∼8,400 years at 20 °C), the image decoded from the DNA-MOF released DNA library maintained high integrity while the unprotected library exhibited progressive information loss. Quantitative analysis of the sequencing results further confirmed the slowed degradation of DNA-MOF during thermal aging (Fig. 6a). After 2 days of heating, the mean PCPM of unprotected library declined to 10.9% of its initial level, while the DNA-MOF sample maintained 39.0% of the original value, confirming the effective protection of DNA-MOFs. As a result, the image decoded from the DNA-MOF released DNA library maintained more similarities than the unprotected samples. For 7 days heating, the DNA-MOF stored image still has a similarity of 99.80% with the original, whereas the unprotected DNA sample retained only 18.31% (Fig. 6b).

**Fig. 6.**
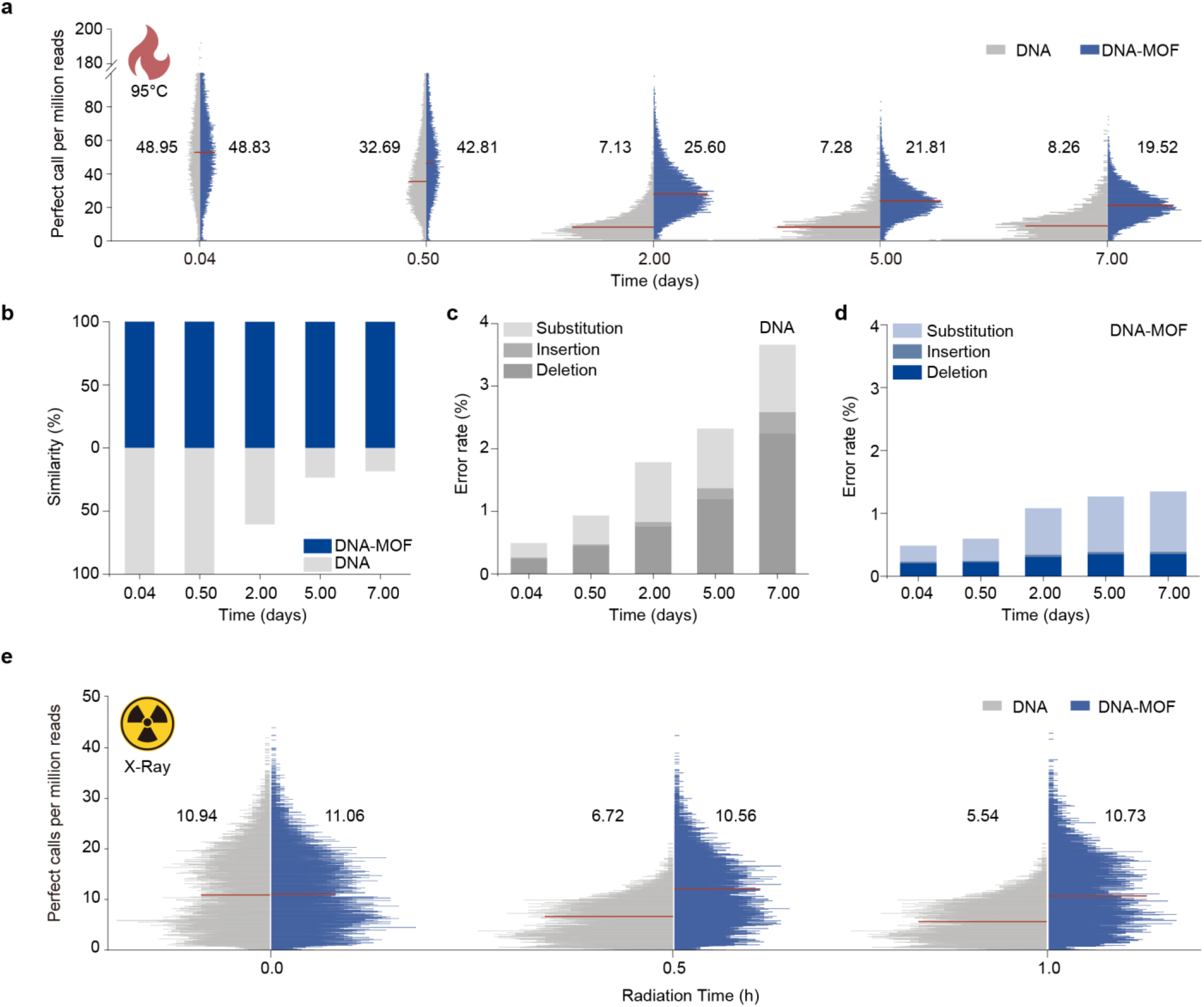
Information integrity evaluation under accelerated aging conditions. (a) Frequency distribution of PCPM for unprotected DNA libraries and DNA-MOF stored libraries after thermal aging. (b) Similarity between the recovered and original images from unprotected DNA libraries and DNA-MOF stored libraries. (c) Error rates per sequence of thermally aged unprotected DNA libraries. (d) Error rates per sequence of thermally aged DNA-MOF stored libraries. (e) Frequency distribution of PCPM for unprotected and DNA-MOF stored libraries after exposure to X-ray irradiation.

We further analyzed inner-strand error rates of covered sequences in the aged libraries. In the unprotected sample, the average deletion error rate increased 2.0% and substitution errors rose 0.80% per sequence after 7 days of aging (Fig. 6c). In contrast, the DNA library released from DNA-MOFs exhibited only a 0.15% increase in deletion errors and a 0.68% increase in substitution errors per sequence (Fig. 6d). All three types of errors (deletions, insertions, and substitutions) were significantly reduced in the DNA-MOF group, confirming its protective role against thermally induced strand breaks and base mutations. Based on the image decoded results, the DNA-MOFs maintained >99% information fidelity even after the equivalent time of four half-lives at 95 °C. This corresponds to an estimated lifespan exceeding 9 millennia at room temperature (20 °C). These results highlight that DNA-MOF can substantially enhance both the fidelity and longevity of DNA-based data storage systems, offering a promising solution to the long-standing challenge of room-temperature stable archival storage.

We further explored the potential of DNA-MOF for extraterrestrial data storage. The unprotected DNA libraries and DNA-MOF samples were exposed to high-intensity X-ray irradiation (10 MGy/h). After 1 h of exposure, the mean PCPM of the unprotected DNA decreased by 49.4%, whereas that of the DNA-MOF sample dropped by only 0.3% (Fig. 6e). This suggests that DNA-MOFs are capable of withstanding extreme irradiation events such as X-ray bursts and may be suitable for data storage in non-terrestrial environments.

## Conclusion

In this work, we proposed a strategy of interweaving DNA with MOF, leading to the formation of DNA-MOFs with high loading density and enhanced protective capabilities. Data stored in DNA-MOFs exhibits strong resistance to various environmental stresses including mechanical, thermal, photonic, and enzymatic degradation. Leveraging these advantages, we established a digital data storage workflow using DNA-MOF as the information medium, which not only matches the storage lifespan of silica-coated DNA, but also offers a high storage density (∼30 wt%) that exceeds the capabilities of conventional protective materials.

Distinct from conventional systems that rely on co-precipitation, adsorption or core-shell structures, DNA-MOFs enable uniformly encapsulation of DNA within the crystal framework, yielding superior DNA loading capacity and theoretical information density up to exabytes per gram. The interwoven crystal structure offers both physical isolation and chemical protection, ensuring reliable data preservation for potentially 9 millennia under room temperature. The lattice framework resembles the zeolites, granting excellent thermal stability of the MOF structure. Furthermore, the crystalline structure provides a physical barrier that resists damage from mechanical shocks and electromagnetic radiation. The pore size distribution characteristics also prevents enzymatic degradation by sterically excluding macromolecular nucleases such as DNase I.

For the convenience of fidelity data integrity assessment, we employed an error-correction-free encoding scheme. DNA-MOF does not impose additional constraints on data encoding, allowing practical implementation with error-correcting codes to repair minor strand-level damage and achieve error-free data recovery over long timescales. Critically, DNA-MOF operates under ambient conditions without specialized cooling or containment, they hold great promise as a foundational material for large-scale cold data storage, potentially reducing the energy demands of cooling systems and network infrastructures in traditional data centers. Additionally, the radiation-shielding capacity of DNA-MOF opens new possibilities for long-term information storage in extraterrestrial environments. Beyond digital data storage, the MOF-biomolecule interweaving strategy can be extended to RNA, proteins and other therapeutic agents, facilitating the design of stable MOF-based carriers for targeted drug delivery and stabilized transport, offering new solutions for handling unstable biomolecules in pharmaceutical applications.

## Notes

### Competing Interest Statement

The authors have declared no competing interest.

### Funding

- National Key R&D Program of China: 2023YFA0915200
- National Natural Science Foundation of China: T2188102, 22322704, U24A20497

